# Chemoproteomic elucidation of β-lactam drug targets in *Mycobacterium abscessus*

**DOI:** 10.64898/2025.12.15.694292

**Authors:** Kaylyn L. Devlin, Emily Hutchinson, Damon T. Leach, Leo J. Gorham, William C. Nelson, Gyanu Lamichhane, Vivian S. Lin, Kimberly E. Beatty

## Abstract

The pathogen *Mycobacterium abscessus* (*Mab*) can cause severe and difficult to treat chronic lung infections. Despite the rising incidence and clinical concern of *Mab* infections, treatment options are limited and often ineffective. Treatment is complicated by *Mab*’s ability to persist in a non-replicative, drug-resistant state. Several β-lactam antibiotics are potently bactericidal against *Mab* but are underutilized because their molecular mechanisms of action against *Mab* are incompletely understood. In the current study, we used β-lactam-derived activity-based probes and chemoproteomics to report the first comprehensive list of enzymes in *Mab* targeted by β-lactams. We compared β-lactam targets across two *Mab* subspecies in actively replicating and non-replicative cultures, using a new carbon starvation model of persistence. We identified 17 targets that were active in every condition tested, seven of which were previously unknown to bind β-lactams. Lastly, we characterized the β-lactamase activity and β-lactam inhibition profiles of nine *Mab* enzymes, demonstrating that imipenem inhibits these targets more effectively than cefoxitin. These findings provide clarity on the mechanisms of action of clinically relevant β-lactams in *Mab*, a crucial step toward fully realizing their potential for treating infections caused by this opportunistic pathogen.

## Introduction

Non-tuberculous mycobacteria (NTM) are a broad group of primarily environmental and non-pathogenic mycobacterial species. However, several NTM cause opportunistic infections of high clinical relevance due to a rapid increase in incidence and antibiotic resistance over recent decades^1^. *Mycobacterium abscessus* (*Mab*) is one of the most significant pathogenic NTM, as it is a top etiological agent of NTM-caused disease^2^.

*Mab*-pulmonary disease (*Mab*-PD) most often develops in patients with existing structural lung disease—such as cystic fibrosis, bronchiectasis, and chronic obstructive pulmonary disease—and is associated with rapid lung function decline and poor clinical outcomes in patients^3^.

*Mab* infections are extremely difficult to cure. There are many innate genetic and physiological traits of *Mab* that make it insensitive to most antibiotics, including high degrees of intrinsic drug resistance, a complex and impermeable cell wall structure, and the expression of drug efflux pumps^4^. Additionally, mycobacteria are known to transition between physiological states in response to microenvironmental conditions within the host, an ability that greatly influences pathogenic susceptibility and survival. *Mab* infects and replicates within macrophages in the lung, leading to the development of necrotic granulomas and survival within caseum^5,6^. These niches expose the pathogen to various physiological conditions and stressors that can include acidic pH, hypoxia, reactive oxygen stress, high lipid concentrations, and/or nutrient deprivation^5,7^. *Mab* is known to respond to these stressors by entering a non-replicative state in which bacilli retain viability without active growth^8–10^. Considering many drugs target bacterial replication and growth, *Mab*’s ability to persist in a non-replicative state is thought to compound drug insusceptibility through phenotypic resistance.

*Mab* drug resistance necessitates long and intensive treatment regimens. The current guidelines for *Mab*-PD treatment involve an intensive phase including an oral macrolide, amikacin, plus an intravenous β-lactam and/or tigecycline^11^. Intensive treatment is followed by a continuation phase involving daily dosing for at least a year, constituting an immense burden on patients. Even with prolonged treatment regimens, *Mab*-PD cure rates are low, ranging from 34-57% depending on *Mab* subspecies^12,13^. Non-replicative *Mab* are thought to contribute to these low cure rates due to decreased susceptibility to most frontline drugs^8,10,14,15^. Clearly, better treatment regimens that are efficacious across *Mab* physiological states are urgently needed.

Compared to other antibiotic classes, there is emerging evidence showing that β-lactams are drivers of overall treatment efficacy^12,16^. Currently, one of two β-lactams is routinely used: either the carbapenem imipenem (IPM) or the cephalosporin cefoxitin (FOX). The choice between β-lactams is made *ad hoc* through antibiotic susceptibility testing^17^, although *in vitro* measurements often fail to correlate with *in vivo* potentcy^5,18^. Better treatment choices could be made if the relative mechanisms of action of β-lactams in *Mab* were more completely understood.

β-lactams inhibit cell wall peptidoglycan biosynthesis^19,20^. In most bacteria, the targets of β-lactams are the penicillin binding proteins (PBPs), which include D,D-transpeptidases (DDTs) and carboxypeptidases. Additionally in mycobacteria, the carbapenem and penem subclasses of β-lactams inhibit a unique class of cell wall crosslinking enzymes known as L,D-transpeptidases (LDTs)^21–27^. However, the targets of β-lactams in *Mab* are much less understood than in other mycobacterial species such as *Mycobacterium tuberculosis*, and a comprehensive list of targets in *Mab* remains to be defined.

The use of β-lactams against *Mab* is complicated by the expression of the extremely active broad-spectrum β-lactamase, Bla_Mab_ (locus identity: MAB_2875), which rapidly hydrolyzes β-lactams^28,29^ and is strongly induced *in vivo*^16^. However, Bla_Mab_ can be inhibited by select β-lactamase inhibitors^30^ and β-lactams used in combination with β-lactamase inhibitors have shown strong potency and synergy against *Mab* both *in vitro*^31–34^ and *in vivo*^16,35,36^. β-lactamase inhibitors are rarely prescribed for *Mab* infections, therefore the optimal use of β-lactams in treating *Mab* infections remains largely untapped^23^.

To effectively utilize β-lactams to treat *Mab*-PD, their targets in *Mab* need to be defined and understood. A powerful approach for such studies is to use activity-based probes (ABPs), which are enzyme substrate analogues that can be used to covalently label and identify drug targets^37^. For example, a green-fluorescent penam, Bocillin FL, has been used in many studies to characterize bacterial enzymes inhibited by β-lactams^38–43^. A limitation of Bocillin FL is that it is highly susceptible to hydrolysis by β-lactamases^28^, such as Bla_Mab_. Additionally, penams have little to no activity against LDTs^21,25,44^, rendering Bocillin FL ineffective at labeling key mycobacterial enzymes integral to cell wall synthesis. By comparison, carbapenems are among the most bactericidal β-lactams against mycobacteria^23,45,46^, are less susceptible to β-lactamase hydrolysis^28,29^, and covalently inhibit LDTs^21,25,26,34,47^. For these reasons, we developed carbapenem-based probes to study β-lactam targets in mycobacteria^22,44^. We demonstrated that a red-fluorescent meropenem ABP is superior to Bocillin FL for studying the regulation and drug susceptibility of β-lactam targets in replicating and hypoxic cultures of *Mycobacterium tuberculosis* (*Mtb*)^44^. Most recently, we used a biotinylated meropenem to comprehensively survey β-lactam targets in *Mtb*^22^. We found that β-lactams target at least 30 enzymes in *Mtb*, indicating a more complex mechanism of action in this pathogen than previously acknowledged.

In the current work, we used ABPs to identify and verify the targets of β-lactams in *Mab* under replicative and non-replicative physiological states. We included two *Mab* strains of different subspecies in our studies: the lab strain ATCC 19977 (subsp. *abscessus*) and the clinical isolate M9510^48^ (subsp. *massiliense*), as these are the dominant subspecies seen in patients. We described a new method for modeling *Mab* persistence by inducing a non-replicative but viable state through carbon starvation. We compared active β-lactam targets in protein gel-resolved lysates from the two strains grown under replicating and carbon-starved conditions. Then, we used meropenem-biotin to comprehensively identify carbapenem targets in *Mab* by activity-based protein profiling (ABPP), a chemoproteomic approach that allows enrichment and identification of drug targets based on enzymatic activity. Lastly, we validated nine *Mab* drug targets through biochemical and drug-binding assays. Our study is the first report of not only a comprehensive list of β-lactam targets in *Mab* but also how these targets are regulated in a clinically relevant persistent state.

## Results and Discussion

### Modeling *Mab* persistence through carbon starvation

A *Mab*-induced lesion in a patient presents the pathogen with many microenvironmental stressors that differentiate *in vivo Mab* growth from patterns traditionally seen in the laboratory. Modeling these conditions *in vitro* has shown that *Mab* can survive in a non-replicating state in response to specific stressors, such as hypoxia and nutrient starvation^8–10^. Studying drug targets in *Mab* under both active replication and a clinically relevant non-replicating state is an important factor in understanding drug efficacy.

Several studies have used complete nutrient starvation (NS) by culture in phosphate buffered saline (PBS) to induce a non-replicative state in *Mab*^8,10,14^. When we cultured *Mab* 19977 under NS, we observed greatly reduced growth relative to standard high nutrient conditions (REP) (**Figure S1**). Cell counts decreased in the first 48 hrs, followed by steady persistence through the end of measurement at 192 hrs (8 days). It was unclear how the initial cell death would impact proteomic analysis, so we used a form of nutrient starvation that restricts carbon sources while maintaining inorganic compounds (carbon starvation, CS, **Table S1**). This model has previously been used to study persistence in *Mtb*^22,49,50^. Similar to NS, *Mab* cell growth under CS was greatly reduced within the first 48 hrs relative to REP. By 48 hrs, CS cultures had reached a non-replicating state, with no change in cell numbers. In contrast, rapid cell division was observed for growth under high nutrient conditions, where cells replicated through 72 hrs before entering stationary phase at numbers almost three orders of magnitude greater than those measured under CS. Carbon starvation was therefore shown to be a simple and reproducible method to quickly induce a non-replicative state to model *Mab* persistence. We used CS at the 72 hr time point in subsequent studies.

### Comparison of β-lactam targets using fluorescent probes

*Mab* strains show variable drug susceptibility across subspecies^51^. To account for this variability and assess potential differences in response to CS across subspecies, we included two strains in our study: a laboratory reference strain of *Mab* (ATCC 19977, subsp. *abscessus*) and a clinical strain obtained from a patient with cystic fibrosis^52^ (M9510, subsp. *massiliense*). We initially evaluated targets of β-lactams in these strains grown under REP and CS conditions using fluorescent ABPs. We compared our published red-fluorescent carbapenem probe, meropenem-sulfoCy5^22^ (Mero-sCy5), to green-fluorescent Bocillin FL. While PBPs are inhibited by most β-lactams, LDTs are primarily inhibited by carbapenems and penems. Mero-sCy5 was therefore expected to label more targets (i.e., PBPs plus LDTs) in *Mab* (**Figure 1A**), while Bocillin FL was expected to solely label PBPs (**Figure 1B**). Indeed, when we examined fluorescent labeling of *Mab* by these probes by super-resolution microscopy (**Figure 1C, Figure S2**), we observed areas of concordant and discordant labeling. While most areas of the cell wall were labeled by both probes (areas of white in channel merge), some regions were primarily bound by Mero-sCy5. This result confirms that the targets of these two subclasses of β-lactam overlap but are not identical in living cells. Additionally, our results suggest that the *Mab* peptidoglycan biosynthetic machinery is not uniformly distributed. Patterns of discordant labeling are consistent with studies in other mycobacterial species, which showed distinct localization of LDT and DDT activity^53,54^.

**Figure 1.**
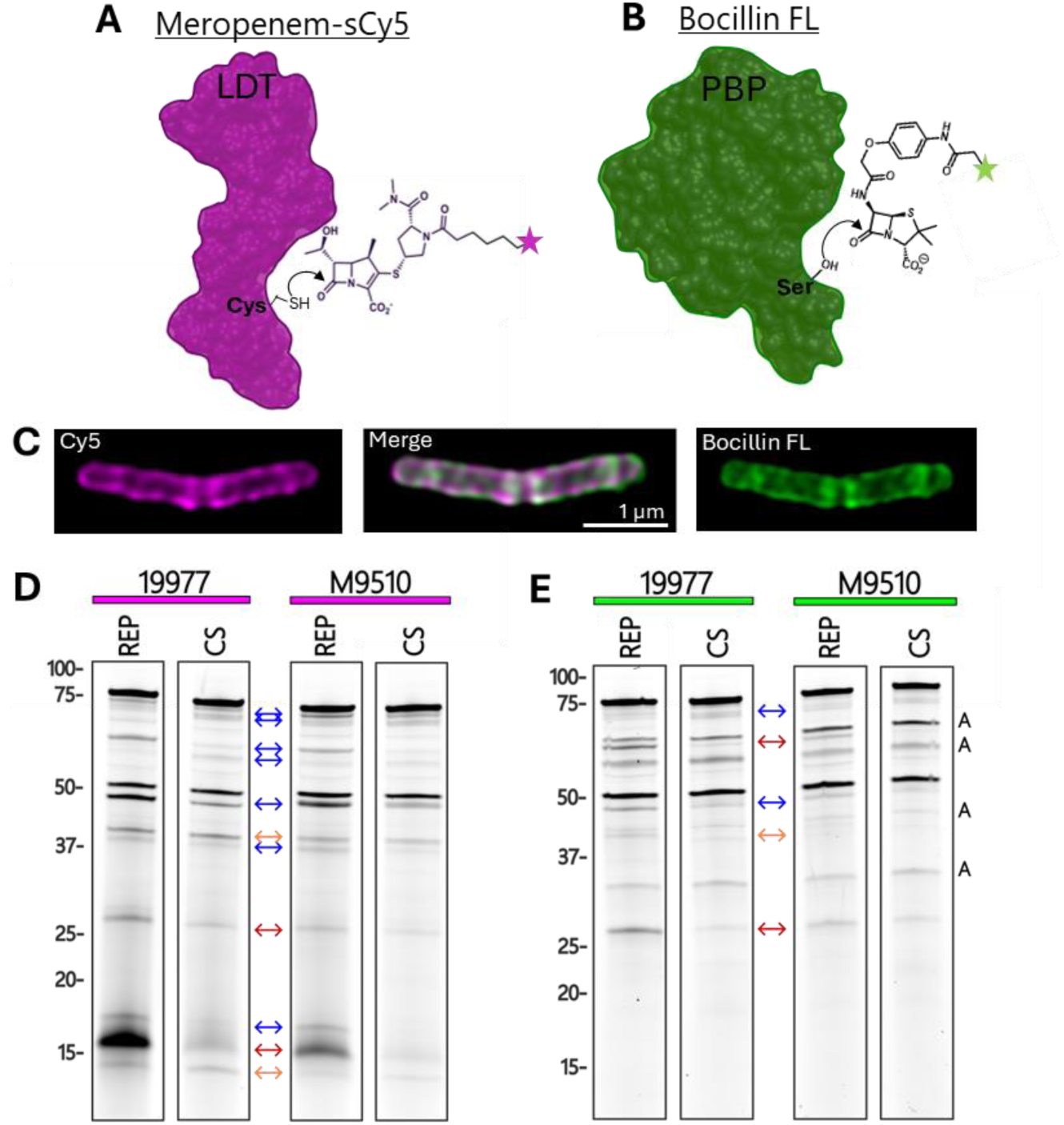
Fluorescent β-lactam probes label protein targets to varying degrees and identify differential activity across *Mab* strains and growth conditions. **A)** The carbapenem probe Meropenem-sCy5 (Mero-sCy5) labels LDTs through reaction with the active-site cysteine. **B)** The pencillin-based probe Bocillin FL labels PBPs, but not LDTs, through reaction with an active-site serine. **C)** Super resolution micrographs of an *Mab* 19977 cell labeled with Mero-sCy5 (Cy5) and Bocillin FL. The middle panel displays a merge of Cy5 and Cy2 channels. **D)** Mero-sCy5-and **E)** Bocillin FL-labeled 19977 and M9510 lysates (30 μg) were resolved by SDS-PAGE (12 μg/lane) and scanned for fluorescent signal. Bands that vary in intensity across groups are marked as follows: between strains within growth conditions (orange arrow), between growth conditions within strains (blue arrow), between both strains and growth conditions (red arrow). Autofluorescent bands in the Cy2 channel (E) are marked with a black A.

We also compared enzyme labeling by these probes in whole-cell lysates from *Mab* 19977 and M9510 cultured in REP and CS conditions (**Figure 1D,E**). Normalized ABP-labeled lysates were resolved by SDS-PAGE and gels were scanned for fluorescent signal (**Figure S3**). Mero-sCy5 (**Figure 1D**) labeled more targets than Bocillin FL (**Figure 1E**), supporting that it binds more enzymes, such as the LDTs. Therefore, a carbapenem ABP outperforms Bocillin FL for labeling β-lactam targets in *Mab* and other mycobacteria^44^.

This analysis also allowed comparison of the activity of β-lactam targets across strains and growth states, as the intensity of labeling corresponds to target abundance, active-site accessibility, and reactivity. We observed numerous differences in target labeling across conditions. Labeling of most targets was weaker in CS than REP. Of note, a faint band around 72 kDa appeared more intensely labeled by both probes in CS relative to REP, suggesting that this unidentified enzyme gains activity and/or abundance in a non-replicating state. While overall target patterns were similar between the strains, several targets appeared less active in M9510 relative to 19977. This finding could suggest a decreased efficacy of β-lactams in M9510. Indeed, past characterization of M9510 (subsp. *massiliense*) identified lower susceptibility to β-lactams relative to 19977 (subsp. *abscessus*)^52^, as evidenced by an increased minimum inhibitory concentration (MIC) for two carbapenems (2.5-fold for IPM and 4-fold for doripenem). Overall, these results demonstrate that ABPs can be used to compare drug targets across strains and growth conditions.

### Mero-biotin activity-based protein profiling

ABPP couples ABP labeling of enzyme targets with mass spectrometry to enable the enrichment and identification of proteins targeted by a molecule of interest. We previously generated a biotinylated version of the meropenem probe (Mero-biotin, **Figure S4**) for ABPP in *Mtb*^22^. In the present study, we used Mero-biotin with ABPP to comprehensively assess the targets of carbapenems in two strains of *Mab* under replicating and carbon-starved conditions. Whole-cell lysates were labeled with Mero-biotin, affinity purified using streptavidin beads, and analyzed by liquid chromatography tandem mass spectrometry (LC-MS/MS). Targets of Mero-biotin were stringently defined based on significantly higher mean intensity values (≥ 3-fold, p ≤ 0.05) relative to mock-labeled control samples. Comprehensive lists of identified targets are included in **Table S2**. Endogenously biotinylated proteins (**Table S3**) were manually removed as false hits.

The final number of Mero-biotin targets identified in the groups analyzed were as follows: 19977 REP (75), 19977 CS (92), M9510 REP (53), M9510 CS (50). Similar to our findings in *Mtb*^22^, these lists include many proteins beyond the PBPs and LDTs that are routinely described as the sole targets of β-lactams. The breadth of targets identified in *Mab* highlights the promiscuous reactivity of these drugs and suggests that polypharmacology is an important feature of β-lactam efficacy across mycobacterial species. A limitation of performing ABPP on whole-cell lysates, however, is the loss of target localization information. We acknowledge that meropenem is unlikely to inhibit all of the identified proteins *in vivo* due to limits on drug penetration or protein accessibility. We plan to address this limitation in future work by conducting ABP labeling in live cells with drug competition. This approach will enable us to expand on the findings presented here by identifying which β-lactam targets are most clinically relevant.

### Targets of carbapenems

A comparison of identified targets across subspecies and growth states highlighted a set of 17 proteins as key carbapenem targets (**Figure 2A**). Identified in all four groups analyzed, this set of key targets is comprised of protein classes known to interact with β-lactams across bacterial species, including D,D-transpeptidases, carboxypeptidases, LDTs, and the broad-spectrum β-lactamase Bla_Mab_. Additionally, five uncharacterized proteins were identified in this central group including LpqF (MAB_0552c), MAB_0330, MAB_2833, MAB_4800, and MAB_4299c. Most of these were bioinformatically annotated as probable β-lactamases in Interpro^55^ (**Figure 2B**, IPR012338) but were not experimentally validated. Importantly, all 17 of these proteins are expressed, active, and inhibited by carbapenems in two different *Mab* subspecies under carbon-deprived conditions modeling granuloma microenvironments.

**Figure 2.**
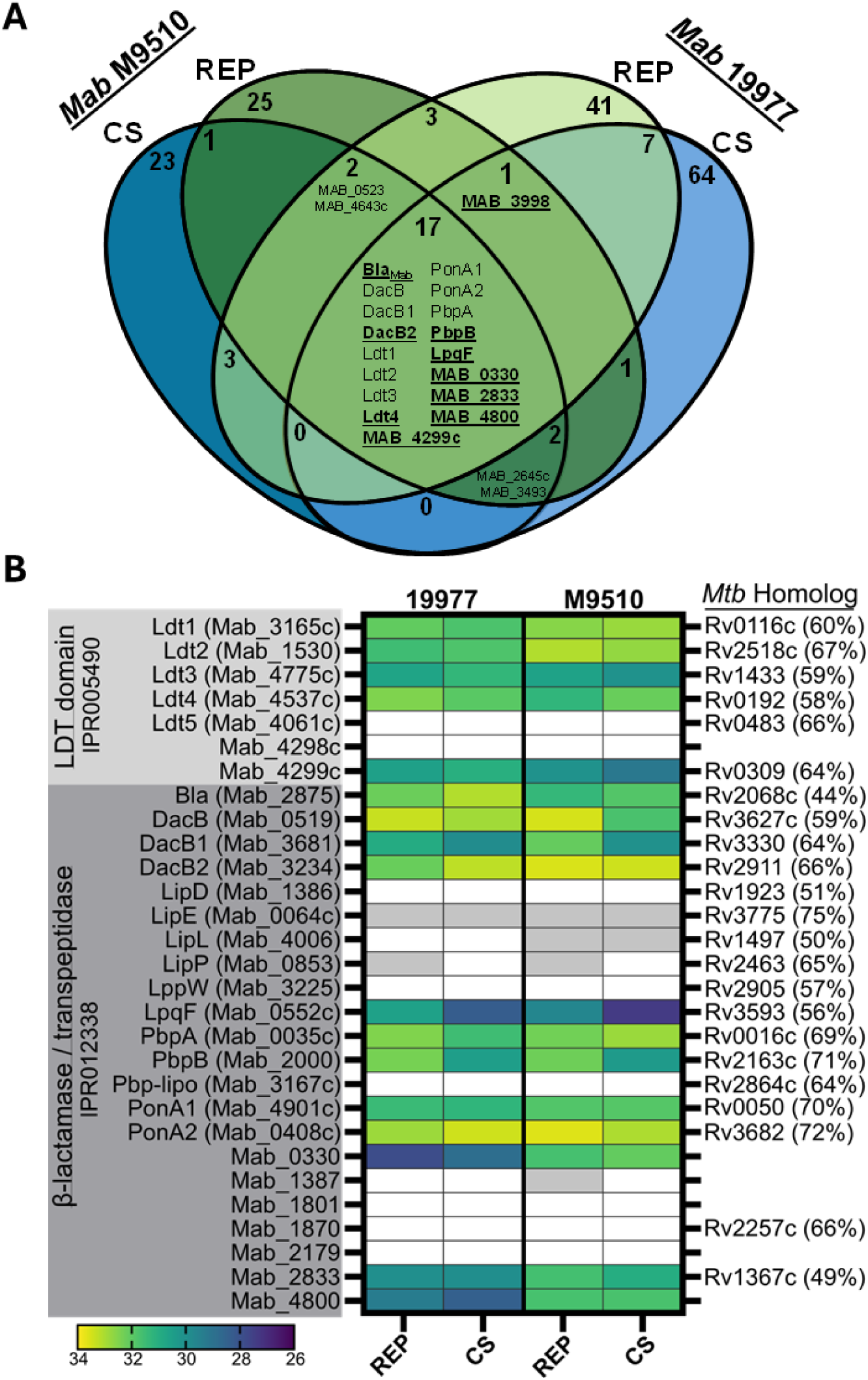
Mero-biotin targets identified by ABPP. **A)** Venn diagram of shared Mero-biotin target proteins identified in *Mab* 19977 and *Mab* M9510 exposed to nutrient rich (REP) and carbon starved (CS) growth conditions. Proteins highlighted in underlined boldface text were selected for further characterization. **B)** Heat map of *Mab* proteins with an Interpro protein family classification of LDT catalytic domain (IPR005490) or β-lactamase/transpeptidase-like (IPR012338). Heat map displays mean log2 label-free quantitation intensity. A white box indicates a protein was found in n < 3 samples. A grey box indicates the protein was found in that group (n ≥ 3) but was not identified as a Mero-biotin hit after stringent filtering.

PBPs and LDTs are well documented to interact with carbapenems in *Mtb*, but these interactions have only recently been investigated in *Mab*. The D,D-carboxypeptidase DacB2 (MAB_3234)^34,47^ was recently validated to bind carbapenems, a finding supported by our results. Similarly, recent evidence showed that PonA1, PonA2, PbpA, and PbpB bind to β-lactams to varying degrees, with carbapenems displaying the most potent inhibition^34,38,47^. The present results are the first validation that the carboxypeptidases DacB (MAB_0519) and DacB1 (MAB_3681) bind β-lactams in *Mab*. This is also the first report of β-lactam binding by the predicted β-lactamases LpqF, MAB_0330, MAB_2833, and MAB_4800. These enzymes have a conserved active site motif (Ser-X-X-Lys) characteristic of a Class A β-lactamase.

There were seven *Mab* proteins classified as containing the L,D-transpeptidase catalytic domain in Interpro^55^ (**Figure 2B**, IPR005490). We identified five of these as carbapenem targets in both strains analyzed: Ldt1, Ldt2, Ldt3, Ldt4, and MAB_4299c.

These results are in accordance with our findings in *Mtb*^22^, where we identified the homologs of each of these enzymes. Our results in lysates support previous reports of recombinant *Mab* Ldt1^25^, Ldt2^25^, Ldt3^34,47^, and Ldt4^34,38,47^ binding carbapenems.

Identification of MAB_4299c, a homolog of *Mtb* protein Rv0309 (64% sequence identity), is interesting because both enzymes have an LDT active site with a conserved cysteine. In prior work, we confirmed that this cysteine is involved in the binding of Rv0309 with β-lactams^22^. Furthermore, Rv0309 localizes to the cell wall, where it decreases cell permeability and enhances mycobacterial survival inside macrophages^56^. These findings in *Mtb* suggest beneficial effects of inhibiting MAB_4299c in *Mab*. We did not identify Ldt5 in the present study, in agreement with previous reports that found no interaction between *Mab* Ldt5 and most β-lactams^34,47^ except the penem sulopenem^57^.

Overall, our chemoproteomic results define a comprehensive list of β-lactam targets in replicating and non-replicating *Mab*. Our data further support the interaction of carbapenems with PBPs and LDTs, while providing the first evidence of β-lactam binding to DacB, DacB1, LpqF, MAB_4299c, MAB_0330, MAB_2833, and MAB_4800. Although our ABPP approach does not provide insight into which are the most clinically relevant targets during infection, the inhibition of many of these proteins is expected to contribute to overall β-lactam efficacy in *Mab*. Notably, PbpB is essential for *Mab* growth *in vitro*, while PonA1, PonA2, DacB2, Ldt1, and Ldt2 confer a growth advantage^58^.

### Differences in target enrichment in CS

All identified carboxypeptidases and transpeptidases were found in both replicating and non-replicating conditions, although some were differentially enriched (**Figure 2B**).

These variations could be due to differences in protein abundance or activity, leading to altered interactions with Mero-biotin. In 19977, there were five differentially enriched targets. PbpB (3.7-fold, p = 0.0001), LpqF (4.7-fold, p = 1.3e-5), and DacB1(2-fold, p = 0.0007) were more enriched in REP than CS. Bla_Mab_ (1.9-fold, p = 0.03) and PonA2 (1.5-fold, p = 0.01) were more enriched in CS.

Most of these targets were also differentially enriched in M9510. PbpB (3.8-fold, p = 2.5e-5), LpqF (4.9-fold, p = 0.0009), and DacB1 (4.3-fold, p = 1.9e-6) were similarly more enriched in REP than CS. There were also some notable differences. DacB (3.5-fold, p = 3.8e-7) and PonA2 (1.5-fold, p = 0.02) were more enriched in M9510 REP. Only Ldt4 was significantly more enriched in CS compared to REP (1.9-fold, p = 0.03). While these variations were mostly congruent between the two *Mab* strains, the directionality of PonA2 differential enrichment was opposite across 19977 and M9510.

### Characterization of target β-lactamase activity

We selected a subset of the key targets for characterization of β-lactamase function, including PbpB, DacB2, Ldt4, LpqF, MAB_0330, MAB_2833, MAB_4299c, and MAB_4800. We also analyzed MAB_3998, which was identified as a carbapenem target in three out of the four proteomic datasets. This protein was of interest due to the presence of two β-lactamase motifs (Ser-X-X-Lys) in its sequence.

These nine recombinant proteins were expressed and purified side-by-side under native conditions (**Figure S5**). β-lactamase function was initially assessed using the chromogenic cephalosporin nitrocefin (**Figure 3A**). We observed nitrocefin hydrolysis above buffer-only control from all enzymes. The same analysis with recombinant Bla_Mab_ resulted in complete nitrocefin hydrolysis near instantaneously (data not shown), consistent with published reports^28^. Enzymes that were bioinformatically predicted to be β-lactamases, including MAB_0330, MAB_2833, and MAB_4800, showed the lowest degree of activity. Unexpectedly, the enzymes with the highest β-lactamase activity were Ldt4 and PbpB.

**Figure 3.**
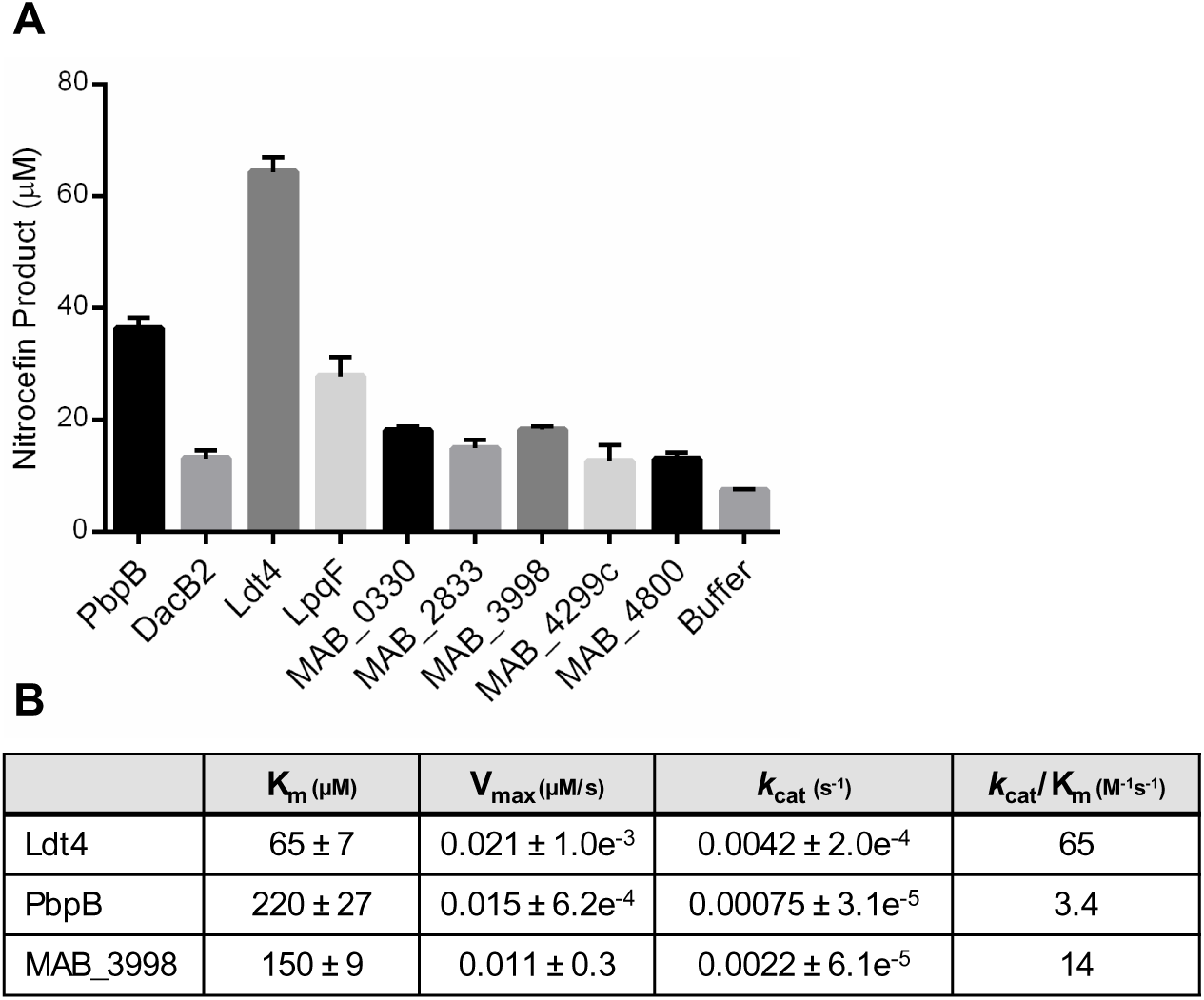
β-lactamase activity of key carbapenem target proteins in *Mab*. **A)** Endpoint measurements of hydrolyzed product following incubation of *Mab* enzymes with nitrocefin. Purified recombinant protein (8 μM) was mixed with nitrocefin (200 μM) and incubated for 2 hr at 37 °C. Absorbance at 486 nm was measured and hydrolyzed product concentrations were determined using Beer’s law. Mean values and standard deviation of technical triplicates were plotted. Data shown is respresentative of two independent experiments. The value of nitrocefin hydrolyzed by each protein was compared to buffer control by multiple unpaired t-tests. Each enzyme produced significantly more hydrolyzed product (p < 0.05). **B)** Steady-state kinetic parameters for nitrocefin hydrolysis by Ldt4, PbpB, and MAB_3998.

To further characterize these findings, we determined the steady-state kinetic parameters for the hydrolysis of nitrocefin by Ldt4, PbpB, and MAB_3998 (**Figure 3B**). Each had micromolar affinity (K_m_) for nitrocefin with low turnover rates (*k*_cat_) and poor catalytic efficiencies (*k*_cat_/K_m_). These results support binding of these enzymes to β-lactams but suggest that their β-lactamase activity is too low to confer drug resistance. Comparatively, Bla_Mab_ has many orders of magnitude greater catalytic efficiency for a wide range of β-lactams, including nitrocefin (*k*_cat_/K_m_ = 4.3e^7^ M^-1^s^-1^)^28,29^. Both Ldt1 and Ldt2 in *Mab* were previously found to have β-lactamase activity, with Ldt2 having a weaker affinity for nitrocefin (328 μM) but a faster maximal rate of hydrolysis (0.33 μM/s) compared to Ldt4^25^.

### Characterization of target inhibition by clinically relevant β-lactams

Next, we used a competition assay to determine how effectively clinically used β-lactams inhibit each of the validated enzymes. We selected IPM and FOX because standard regimens to treat *Mab* infections use these drugs. We also included the β-lactamase inhibitor avibactam (AVI). AVI is not currently considered for use for *Mab* infections but is effective at inhibiting Bla_Mab_^30,32^. We assessed protein inhibition after drug treatment through quantification of fluorescent ABP binding (**Figure 4, Figure S6**). Purified protein was incubated with drug or buffer only before labeling with either Mero-sCy5 or Bocillin FL. Samples were then resolved by SDS-PAGE and imaged for quantification. All proteins were inhibited by IPM, while FOX did not inhibit Ldt4 or the putative LDT MAB_4299c. MAB_0330 was weakly inhibited by FOX. AVI treatment mostly had no effect on ABP binding, except for an intermediate inhibition of DacB2 and an enhancement of Ldt4 binding by Mero-sCy5. The β-lactam preference of several proteins was also evident from varying probe responses. MAB_4800 bound Mero-sCy5 better than Bocillin FL while MAB_0330 was the opposite.

**Figure 4.**
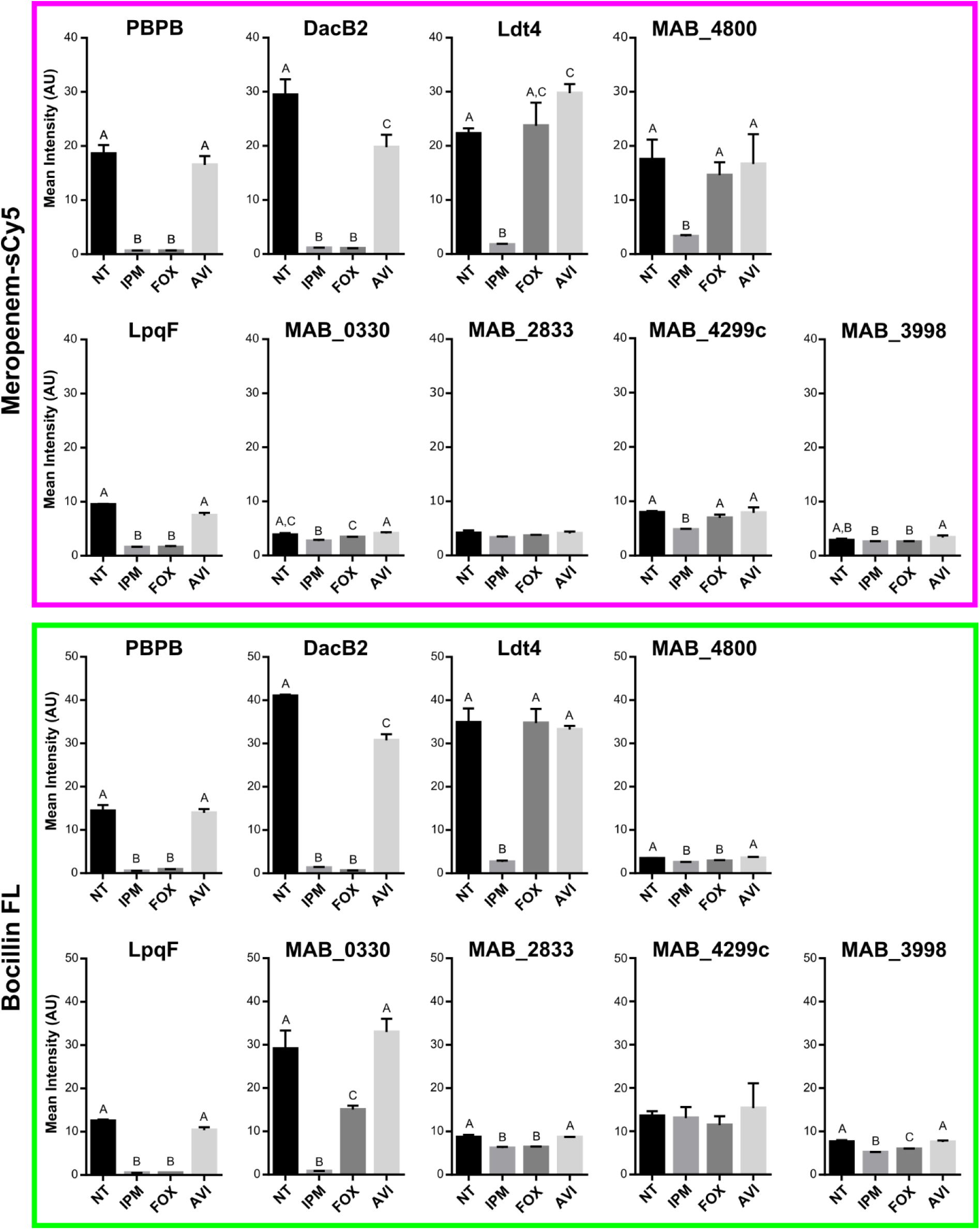
Imipenem inhibits a wide range of β-lactam interacting proteins, while cefoxitin inhibits only a subset. Mean intensity of protein gel-resolved fluorescent bands after treatment with β-lactam drugs. Purified protein (3 μg) was combined with drug (500 μM) or buffer (no treatment, NT) in 10 mM HEPES pH 7.5 (30 min, 37 °C). Samples were then labeled with 10 μM Mero-sCy5 or Bocillin FL (1 hr, 37 °C). Labeled protein was resolved via SDS-PAGE (1 μg/lane) and fluorescent signal was quantified as mean intensity of three independent replicates. Error is reported as standard deviation and statistical significance was determined by one-way ANOVA (α = 0.05).

We determined the affinity of Ldt4, PbpB, DacB2, and MAB_4800 for FOX and IPM by measuring the drug concentration that inhibited enzymatic activity by 50% (IC50). We treated each enzyme with a range of drug concentrations before Bocillin FL labeling.

Samples were resolved by SDS-PAGE before quantitative measurement of fluorescent band intensity, a method described by Kocaoglu and Carlson^59^. Dose-response curves were plotted to determine the IC50 for each enzyme (**Figure 5** and **Figure S7**). All enzymes were potently inhibited by IPM at low micromolar doses. FOX did not inhibit Ldt4 or MAB_4800 in the concentration range analyzed. IPM inhibited PbpB 3-fold more potently than FOX. In contrast, FOX inhibited DacB2 5-fold more potently than IPM. These trends matched what we observed qualitatively in **Figure 4**.

**Figure 5.**
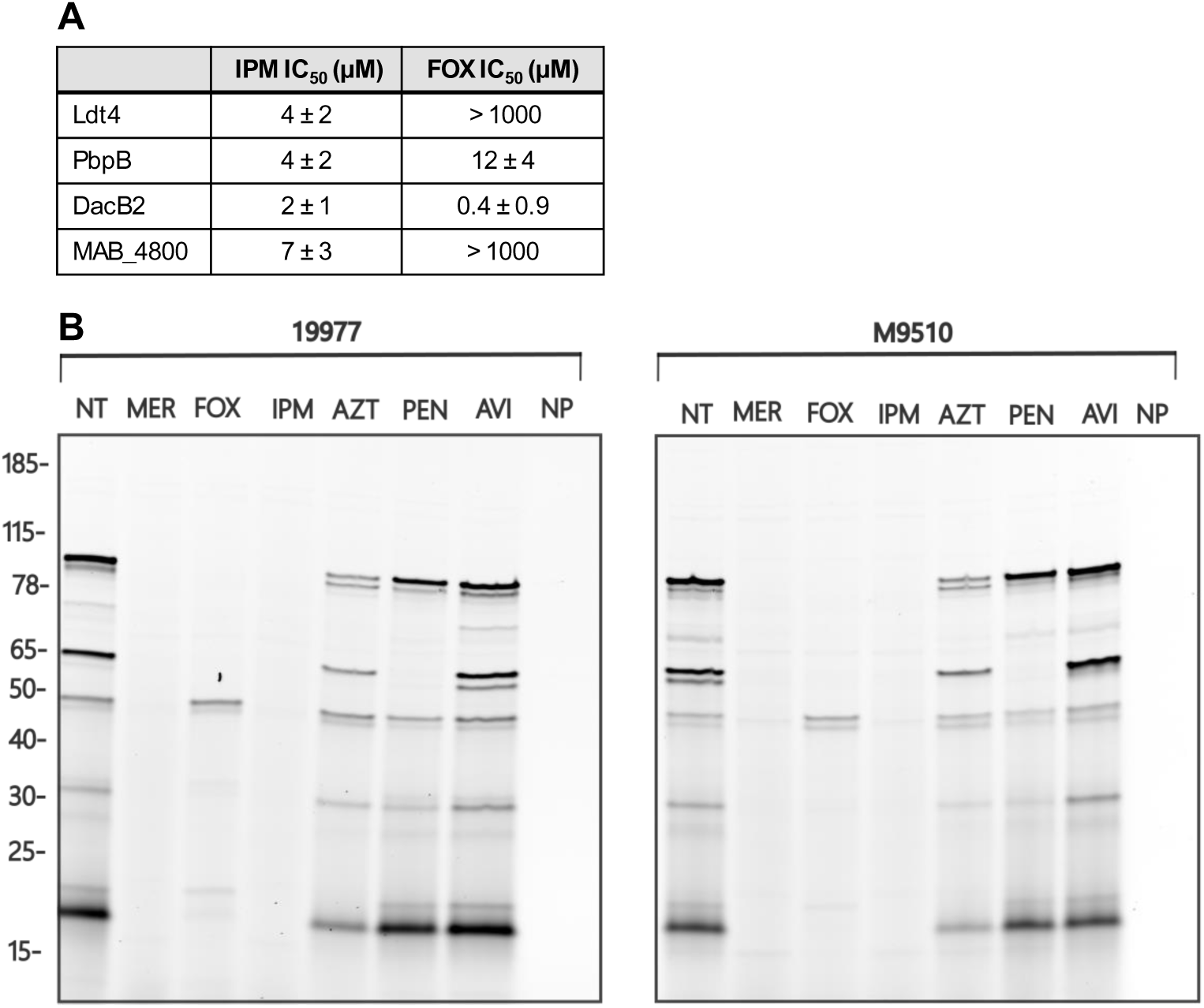
Carbapenems inhibit *Mab* enzymes more potently than other β-lactam subclasses. **A)** Drug IC_50_ values for Ldt4, PbpB, DacB2, and MAB_4800. **B)** Comparison of drug inhibition of β-lactam targets in gel-resolved lysates of *Mab* 19977 and M9510. Lysates (8 μg) were pretreated with drug (1 mM) or buffer only (no treatment, NT) and labeled with Mero-sCy5 (5 μM) or buffer only (no probe, NP). Labeled lysates were resolved by SDS-PAGE (5 μg/lane) and scanned for fluorescent signal. Drugs used were as follows: meropenem (MER), cefoxitin (FOX), imipenem (IPM), aztreonam (AZT), penicillin G (PEN), avibactam (AVI).

We compared these IC50 values to published β-lactam MIC_90_ values and median unbound average steady-state plasma concentrations (*f*C_ss,avg_). Hunkins et al.^51^ reported an MIC_90_ of 16 μg/mL for IPM and 64 μg/mL for FOX in both *M. abscessus* subsp. *abscessus* (n = 1344 isolates) and *M. abscessus* subsp. *massiliense* (n = 754 isolates). Sayed et al.^38^ reported an *f*C_ss,avg_ achievable in 50% of patients receiving clinically relevant drug doses of 12.2 μg/mL for IPM and 4.0 μg/mL for FOX. Both the MIC_90_ and *f*C_ss,avg_ for IPM are above the corresponding IC50 we determined for these four enzymes, which are 0.5-2 μg/mL. This supports that the IPM doses used for treatment of standard *Mab* infections would likely inhibit all four of these enzymes, including the essential enzyme PbpB. On the other hand, FOX would potentially only inhibit DacB2, as its *f*C_ss,avg_ is below the IC50s measured for the other enzymes.

Finally, we compared the relative effectiveness of diverse β-lactams at inhibiting targets in whole cell lysates from 19977 and M9510. We tested two carbapenems (meropenem and IPM), FOX, the monobactam aztreonam (AZT), the penam penicillin G (PEN), and the β-lactamase inhibitor AVI. Lysates were treated with excess drug before labeling with Mero-sCy5 (**Figure 5B**). The carbapenems abolished ABP labeling, indicating that both inhibited all targets. FOX inhibited many targets but not all. AZT and PEN were largely ineffective inhibitors, apart from few responsive proteins. Together, these data definitively demonstrated that IPM is more effective than other β-lactams, including FOX, at inhibiting β-lactam target enzymes in two subspecies of *Mab*.

### Carbapenem target overlap between *M. abscessus* and *M. tuberculosis*

We recently used ABPP to identify targets of β-lactams in *M. tuberculosis* under CS and REP growth conditions^22^. Using those published datasets, we identified target overlap between *M. tuberculosis* and *M. abscessus* (**Table 1**). There were fifteen targets identified in both growth conditions across species. Targets included three D,D-carboxypeptidases, four D,D-transpeptidases, five LDTs, and three β-lactamases.

**Table 1.**
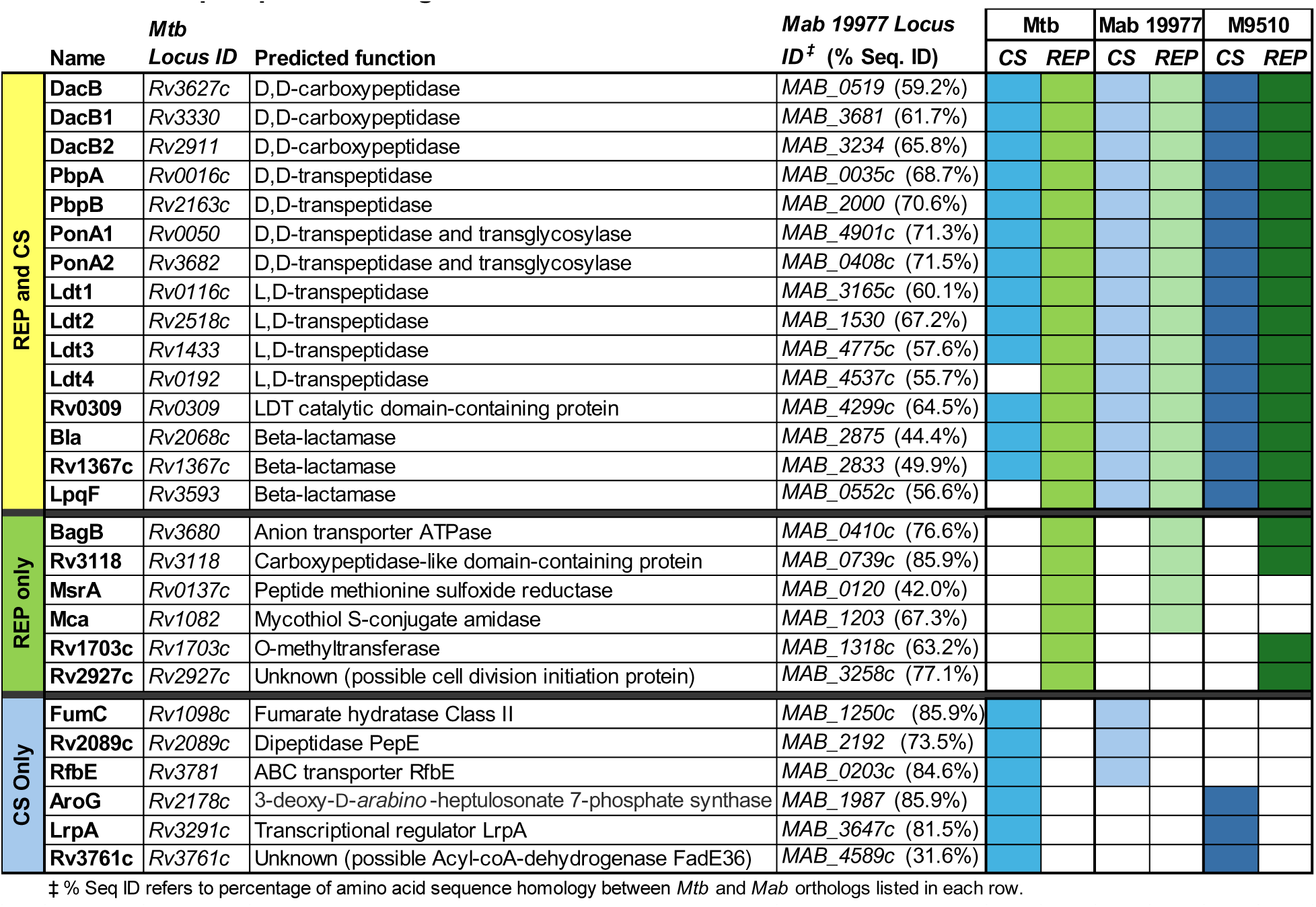
Overlap of β-lactam targets identified in *Mtb* and *Mab* under REP and CS conditions.

Peptidoglycan biosynthesis and structure are well conserved between these two pathogens, so identifying these targets by ABPP in both species was expected. This congruence served to validate the robustness of the identification of β-lactam targets among strains and species.

We examined targets that were identified in both species under REP and CS conditions to speculate on how these proteins might interact with β-lactams (**Table 1**). There were six proteins identified in both species in REP conditions alone. This list included MAB_0739c, a protein of unknown function whose *Mtb* ortholog Rv3118 (85.9% sequence identity) is also uncharacterized. We determined that both MAB_0739c and Rv3118 encode a carboxypeptidase-like domain (IPR008969) found in metallo-carboxypeptidases. We hypothesize that these proteins bind carbapenems through that domain. Next, we considered MAB_3258c, which is annotated in Uniprot^60^ as a possible cell division initiation protein. MAB_3258’s closest *Mtb* ortholog, Rv2927c (77% sequence identity), is essential^61^, but its function and localization are unknown.

MAB_3258c is more distantly related to Wag31 (Rv2145c; 36% sequence identity), a scaffolding protein involved in regulation of cell shape, growth, and cell wall biosynthesis^62,63^. Wag31 has numerous interaction partners, including PbpB^62,64^. We surmise that we identified MAB_3258 and Rv2927c not because either directly bind carbapenems, but rather through tight binding interactions with a carbapenem-bound PBP^65^. An alternative explanation is that these proteins were enriched through interactions with biotinylated acetyl-coA carboxylases, which also bind Wag31^66^.

Six proteins were identified solely in CS in both species. We were intrigued by the function of two of these, AroG and LrpA. Firstly, we identified AroG (MAB_1987 and Rv2178c). AroG is an essential 3-deoxy-D-*arabino*-heptulosonate 7-phosphate synthase that catalyzes the first committed step of aromatic amino acid biosynthesis. This enzyme’s structure and activity is well described in *M. tuberculosis*^67^. Unusually, AroG activity is controlled synergistically by three allosteric sites that bind aromatic amino acids (i.e., Phe, Tyr, and Trp)^68,69^. It is interesting to consider that a β-lactam might bind one of those sites, altering AroG’s catalytic activity. Secondly, we identified MAB_3647c, an ortholog of the transcriptional regulator LrpA (Rv3291c). LrpA is upregulated in nutrient starvation^70^, supporting a key role in persistence. LrpA binding to DNA is modulated by effectors, including aromatic amino acids^71^. We speculate that β-lactams could be a previously unknown effector for LrpA.

## Conclusions

*Mab*-PD is characterized by heterogeneous lesions with variable distribution of actively replicating and non-replicating *Mab*. Susceptibility to β-lactam antibiotics is influenced by the targets expressed in these distinct physiological states. To identify β-lactam targets relevant to each state, we used *in vitro* models of replicating and non-replicating conditions and analyzed two strains representing the *abscessus* and *massiliense* subspecies. We identified 17 carbapenem-binding proteins active in both strains and across both growth states. Notably, 15 of these proteins are also targets in *Mtb*^22^, suggesting that they are conserved mycobacterial drug targets.

β-lactam targets included cell wall biosynthesis enzymes, β-lactamases, and uncharacterized proteins containing a β-lactamase sequence motif. This is the first description of β-lactam binding to seven *Mab* proteins (DacB, DacB1, LpqF, MAB_4299c, MAB_2833, MAB_0330, MAB_4800). Importantly, key targets were shown to be maintained and active in the non-replicating state, which contributes to bacterial drug resistance and survival in a *Mab* infection. This suggests that these enzymes would be inhibited by carbapenems in persistent *Mab* that are no longer responsive to many frontline drugs.

We characterized the β-lactamase activity and drug inhibition profiles of nine enzymes (PbpB, DacB2, Ldt4, LpqF, MAB_4800, MAB_0330, MAB_2833, MAB_4299c, MAB_3998). Ldt4 and PbpB displayed the highest β-lactamase activity, although this activity is orders of magnitude less than Bla_Mab_. We measured the inhibition of these enzymes by frontline β-lactams used to treat *Mab*-PD: IPM and FOX. IPM was superior at inhibiting *Mab* targets, both against purified enzymes and in whole-cell lysates. Our findings define molecular mechanisms contributing to IPM efficacy against *Mab* and suggest that it should be used preferentially over FOX in the clinic. Overall, the current work provides novel insights into the mechanism of action of β-lactams against *Mab*.

## Materials and methods

Detailed experimental protocols are included in the electronic supporting information.

## Acknowledgements

Funding for the Beatty group was provided by the National Institute of Health (NIAID: R01 AI149737 and NIGMS: R35 GM149559). Pacific Northwest National Laboratory is operated by Battelle for the Department of Energy (DOE) under contract DE-AC05-76RL01830. A portion of this research was supported by an Environmental Molecular Sciences Laboratory (EMSL) user project award (https://www.emsl.pnnl.gov/project/60231) for leveraging instrument capabilities operating at EMSL, a DOE Office of Science User Facility. Development of chemoprotR was supported by an EMSL Science and Technology Developer project award (https://www.emsl.pnnl.gov/project/60473). We thank Dr. Jordan Devereux (OHSU Medicinal Chemistry Core; RRID: SCR_019048) for synthesizing Mero-sCy5 and Dr. Felice Kelly (OHSU Advanced Multiscale Microscopy Core; RRID: SCR_009961) for expert technical assistance with super-resolution microscopy. We are grateful to Dr. Priscila Lalli and Dr. Ron Moore for proteomics runs and Dr. Matt Monroe for proteomics data support.

## Citations

1 Victoria, L., Gupta, A., Gomez, J. L. & Robledo, J. Mycobacterium abscessus complex: A Review of Recent Developments in an Emerging Pathogen. Front Cell Infect Microbiol 11, 659997 (2021). 10.3389/fcimb.2021.659997

2 Johansen, M. D., Herrmann, J. L. & Kremer, L. Non-tuberculous mycobacteria and the rise of Mycobacterium abscessus. Nat Rev Microbiol 18, 392–407 (2020). 10.1038/s41579-020-0331-1

3 Harris, K. A. & Kenna, D. T. D. Mycobacterium abscessus infection in cystic fibrosis: molecular typing and clinical outcomes. J Med Microbiol 63, 1241–1246 (2014). 10.1099/jmm.0.077164-0

4 Falkinham, J. O., 3rd. Challenges of NTM Drug Development. Front Microbiol 9, 1613 (2018). 10.3389/fmicb.2018.01613

5 Wu, M. L., Aziz, D. B., Dartois, V. & Dick, T. NTM drug discovery: status, gaps and the way forward. Drug Discov Today 23, 1502–1519 (2018). 10.1016/j.drudis.2018.04.001

6 Kaya, F., Ernest, J. P., LoMauro, K., Gengenbacher, M., Madani, A., Aragaw, W. W., Zimmerman, M. D., Sarathy, J. P., Alvarez, N., Daudelin, I., Wang, H., Lanni, F., Weiner, D. M., Via, L. E., Barry, C. E., 3rd, Olivier, K. N., Dick, T., Podell, B. K., Savic, R. M. & Dartois, V. A Rabbit Model to Study Antibiotic Penetration at the Site of Infection for Nontuberculous Mycobacterial Lung Disease: Macrolide Case Study. Antimicrob Agents Chemother 66, e0221221 (2022). 10.1128/aac.02212-21

7 Sarathy, J. P. & Dartois, V. Caseum: a Niche for Mycobacterium tuberculosis Drug-Tolerant Persisters. Clin Microbiol Rev 33 (2020). 10.1128/CMR.00159-19

8 Yam, Y. K., Alvarez, N., Go, M. L. & Dick, T. Extreme Drug Tolerance of Mycobacterium abscessus "Persisters". Front Microbiol 11, 359 (2020). 10.3389/fmicb.2020.00359

9 Lanni, A., Borroni, E., Iacobino, A., Russo, C., Gentile, L., Fattorini, L. & Giannoni, F. Activity of Drug Combinations against Mycobacterium abscessus Grown in Aerobic and Hypoxic Conditions. Microorganisms 10 (2022). 10.3390/microorganisms10071421

10 Berube, B. J., Castro, L., Russell, D., Ovechkina, Y. & Parish, T. Novel Screen to Assess Bactericidal Activity of Compounds Against Non-replicating Mycobacterium abscessus. Front Microbiol 9, 2417 (2018). 10.3389/fmicb.2018.02417

11 Floto, R. A., Olivier, K. N., Saiman, L., Daley, C. L., Herrmann, J. L., Nick, J. A., Noone, P. G., Bilton, D., Corris, P., Gibson, R. L., Hempstead, S. E., Koetz, K., Sabadosa, K. A., Sermet-Gaudelus, I., Smyth, A. R., van Ingen, J., Wallace, R. J., Winthrop, K. L., Marshall, B. C., Haworth, C. S., Foundation, U. S. C. F. & European Cystic Fibrosis, S. US Cystic Fibrosis Foundation and European Cystic Fibrosis Society consensus recommendations for the management of non-tuberculous mycobacteria in individuals with cystic fibrosis. Thorax 71 **Suppl 1**, i1–22 (2016). 10.1136/thoraxjnl-2015-207360

12 Kwak, N., Dalcolmo, M. P., Daley, C. L., Eather, G., Gayoso, R., Hasegawa, N., Jhun, B. W., Koh, W. J., Namkoong, H., Park, J., Thomson, R., van Ingen, J., Zweijpfenning, S. M. H. & Yim, J. J. Mycobacterium abscessus pulmonary disease: individual patient data meta-analysis. Eur Respir J 54 (2019). 10.1183/13993003.01991-2018

13 Pasipanodya, J. G., Ogbonna, D., Ferro, B. E., Magombedze, G., Srivastava, S., Deshpande, D. & Gumbo, T. Systematic Review and Meta-analyses of the Effect of Chemotherapy on Pulmonary Mycobacterium abscessus Outcomes and Disease Recurrence. Antimicrob Agents Chemother 61 (2017). 10.1128/AAC.01206-17

14 Lee, J., Ammerman, N., Agarwal, A., Naji, M., Li, S. Y. & Nuermberger, E. Differential In Vitro Activities of Individual Drugs and Bedaquiline-Rifabutin Combinations against Actively Multiplying and Nutrient-Starved Mycobacterium abscessus. Antimicrob Agents Chemother 65 (2021). 10.1128/AAC.02179-20

15 Xie, M., Ganapathy, U. S., Lan, T., Osiecki, P., Sarathy, J. P., Dartois, V., Aldrich, C. C. & Dick, T. ADP-ribosylation-resistant rifabutin analogs show improved bactericidal activity against drug-tolerant M. abscessus in caseum surrogate. Antimicrob Agents Chemother 67, e0038123 (2023). 10.1128/aac.00381-23

16 Lefebvre, A. L., Le Moigne, V., Bernut, A., Veckerle, C., Compain, F., Herrmann, J. L., Kremer, L., Arthur, M. & Mainardi, J. L. Inhibition of the beta-Lactamase Bla(Mab) by Avibactam Improves the In Vitro and In Vivo Efficacy of Imipenem against Mycobacterium abscessus. Antimicrob Agents Chemother 61 (2017). 10.1128/AAC.02440-16

17 Daley, C. L., Iaccarino, J. M., Lange, C., Cambau, E., Wallace, R. J., Andrejak, C., Bottger, E. C., Brozek, J., Griffith, D. E., Guglielmetti, L., Huitt, G. A., Knight, S. L., Leitman, P., Marras, T. K., Olivier, K. N., Santin, M., Stout, J. E., Tortoli, E., van Ingen, J., Wagner, D. & Winthrop, K. L. Treatment of Nontuberculous Mycobacterial Pulmonary Disease: An Official ATS/ERS/ESCMID/IDSA Clinical Practice Guideline. Clin Infect Dis 71, 905–913 (2020). 10.1093/cid/ciaa1125

18 Jeon, K., Kwon, O. J., Lee, N. Y., Kim, B. J., Kook, Y. H., Lee, S. H., Park, Y. K., Kim, C. K. & Koh, W. J. Antibiotic treatment of Mycobacterium abscessus lung disease: a retrospective analysis of 65 patients. Am J Respir Crit Care Med 180, 896–902 (2009). 10.1164/rccm.200905-0704OC

19 Hartmann, R., Holtje, J. V. & Schwarz, U. Targets of penicillin action in Escherichia coli. Nature 235, 426–429 (1972). 10.1038/235426a0

20 Mora-Ochomogo, M. & Lohans, C. T. beta-Lactam antibiotic targets and resistance mechanisms: from covalent inhibitors to substrates. RSC Med Chem 12, 1623–1639 (2021). 10.1039/d1md00200g

21 Kumar, P., Kaushik, A., Lloyd, E. P., Li, S. G., Mattoo, R., Ammerman, N. C., Bell, D. T., Perryman, A. L., Zandi, T. A., Ekins, S., Ginell, S. L., Townsend, C. A., Freundlich, J. S. & Lamichhane, G. Non-classical transpeptidases yield insight into new antibacterials. Nat Chem Biol 13, 54–61 (2017). 10.1038/nchembio.2237

22 Devlin, K. L., Hutchinson, E., Dearing, H. N., Levine, S. R., Reid, D. J., Leach, D. T., Griggs, L. H., Lomas, G. X., Gorham, L. J., Wright, A. T., Lamichhane, G., Lin, V. S. & Beatty, K. E. Comprehensive Identification of beta-Lactam Antibiotic Polypharmacology in Mycobacterium tuberculosis. ACS Infect Dis 11, 2422–2433 (2025). 10.1021/acsinfecdis.5c00233

23 Story-Roller, E., Maggioncalda, E. C., Cohen, K. A. & Lamichhane, G. Mycobacterium abscessus and beta-Lactams: Emerging Insights and Potential Opportunities. Front Microbiol 9, 2273 (2018). 10.3389/fmicb.2018.02273

24 Dubee, V., Triboulet, S., Mainardi, J. L., Etheve-Quelquejeu, M., Gutmann, L., Marie, A., Dubost, L., Hugonnet, J. E. & Arthur, M. Inactivation of Mycobacterium tuberculosis l,d-transpeptidase LdtMt(1) by carbapenems and cephalosporins. Antimicrob Agents Chemother 56, 4189–4195 (2012). 10.1128/AAC.00665-12

25 Kumar, P., Chauhan, V., Silva, J. R. A., Lameira, J., d’Andrea, F. B., Li, S. G., Ginell, S. L., Freundlich, J. S., Alves, C. N., Bailey, S., Cohen, K. A. & Lamichhane, G. Mycobacterium abscessus l,d-Transpeptidases Are Susceptible to Inactivation by Carbapenems and Cephalosporins but Not Penicillins. Antimicrob Agents Chemother 61 (2017). 10.1128/AAC.00866-17

26 Cordillot, M., Dubee, V., Triboulet, S., Dubost, L., Marie, A., Hugonnet, J. E., Arthur, M. & Mainardi, J. L. In vitro cross-linking of Mycobacterium tuberculosis peptidoglycan by L,D-transpeptidases and inactivation of these enzymes by carbapenems. Antimicrob Agents Chemother 57, 5940–5945 (2013). 10.1128/AAC.01663-13

27 Bianchet, M. A., Pan, Y. H., Basta, L. A. B., Saavedra, H., Lloyd, E. P., Kumar, P., Mattoo, R., Townsend, C. A. & Lamichhane, G. Structural insight into the inactivation of Mycobacterium tuberculosis non-classical transpeptidase Ldt(Mt2) by biapenem and tebipenem. BMC Biochem 18, 8 (2017). 10.1186/s12858-017-0082-4

28 Soroka, D., Dubee, V., Soulier-Escrihuela, O., Cuinet, G., Hugonnet, J. E., Gutmann, L., Mainardi, J. L. & Arthur, M. Characterization of broad-spectrum Mycobacterium abscessus class A beta-lactamase. J Antimicrob Chemother 69, 691–696 (2014). 10.1093/jac/dkt410

29 Ramirez, A., Ruggiero, M., Aranaga, C., Cataldi, A., Gutkind, G., de Waard, J. H., Araque, M. & Power, P. Biochemical Characterization of beta-Lactamases from Mycobacterium abscessus Complex and Genetic Environment of the beta-Lactamase-Encoding Gene. Microb Drug Resist 23, 294–300 (2017). 10.1089/mdr.2016.0047

30 Soroka, D., Ourghanlian, C., Compain, F., Fichini, M., Dubee, V., Mainardi, J. L., Hugonnet, J. E. & Arthur, M. Inhibition of beta-lactamases of mycobacteria by avibactam and clavulanate. J Antimicrob Chemother 72, 1081–1088 (2017). 10.1093/jac/dkw546

31 Kaushik, A., Ammerman, N. C., Lee, J., Martins, O., Kreiswirth, B. N., Lamichhane, G., Parrish, N. M. & Nuermberger, E. L. In Vitro Activity of the New beta-Lactamase Inhibitors Relebactam and Vaborbactam in Combination with beta-Lactams against Mycobacterium abscessus Complex Clinical Isolates. Antimicrob Agents Chemother 63 (2019). 10.1128/AAC.02623-18

32 Kaushik, A., Gupta, C., Fisher, S., Story-Roller, E., Galanis, C., Parrish, N. & Lamichhane, G. Combinations of avibactam and carbapenems exhibit enhanced potencies against drug-resistant Mycobacterium abscessus. Future Microbiol 12, 473–480 (2017). 10.2217/fmb-2016-0234

33 Dousa, K. M., Kurz, S. G., Taracila, M. A., Bonfield, T., Bethel, C. R., Barnes, M. D., Selvaraju, S., Abdelhamed, A. M., Kreiswirth, B. N., Boom, W. H., Kasperbauer, S. H., Daley, C. L. & Bonomo, R. A. Insights into the l,d-Transpeptidases and d,d-Carboxypeptidase of Mycobacterium abscessus: Ceftaroline, Imipenem, and Novel Diazabicyclooctane Inhibitors. Antimicrob Agents Chemother 64 (2020). 10.1128/AAC.00098-20

34 Shin, E., Dousa, K. M., Taracila, M. A., Bethel, C. R., Nantongo, M., Nguyen, D. C., Akusobi, C., Kurz, S. G., Plummer, M. S., Daley, C. L., Holland, S. M., Rubin, E. J., Bulitta, J. B., Boom, W. H., Kreiswirth, B. N. & Bonomo, R. A. Durlobactam in combination with beta-lactams to combat Mycobacterium abscessus. Antimicrob Agents Chemother 69, e0117424 (2025). 10.1128/aac.01174-24

35 Story-Roller, E., Maggioncalda, E. C. & Lamichhane, G. Select beta-Lactam Combinations Exhibit Synergy against Mycobacterium abscessus In Vitro. Antimicrob Agents Chemother 63 (2019). 10.1128/AAC.02613-18

36 Lefebvre, A. L., Dubee, V., Cortes, M., Dorchene, D., Arthur, M. & Mainardi, J. L. Bactericidal and intracellular activity of beta-lactams against Mycobacterium abscessus. J Antimicrob Chemother 71, 1556–1563 (2016). 10.1093/jac/dkw022

37 Lembke, H. K. & Carlson, E. E. Activity-based probes in pathogenic bacteria: Investigating drug targets and molecule specificity. Curr Opin Chem Biol 76, 102359 (2023). 10.1016/j.cbpa.2023.102359

38 Sayed, A. R. M., Shah, N. R., Basso, K. B., Kamat, M., Jiao, Y., Moya, B., Sutaria, D. S., Lang, Y., Tao, X., Liu, W., Shin, E., Zhou, J., Werkman, C., Louie, A., Drusano, G. L. & Bulitta, J. B. First Penicillin-Binding Protein Occupancy Patterns for 15 beta-Lactams and beta-Lactamase Inhibitors in Mycobacterium abscessus. Antimicrob Agents Chemother 65 (2020). 10.1128/AAC.01956-20

39 Grunnvag, J. S., Hegstad, K. & Lentz, C. S. Activity-based protein profiling of serine hydrolases and penicillin-binding proteins in Enterococcus faecium. FEMS Microbes 5, xtae015 (2024). 10.1093/femsmc/xtae015

40 Gee, K. R., Kang, H. C., Meier, T. I., Zhao, G. & Blaszcak, L. C. Fluorescent Bocillins: synthesis and application in the detection of penicillin-binding proteins. Electrophoresis 22, 960–965 (2001). 10.1002/1522-2683()22:5<960::AID-ELPS960>3.0.CO;2-9

41 Sharifzadeh, S., Dempwolff, F., Kearns, D. B. & Carlson, E. E. Harnessing beta-Lactam Antibiotics for Illumination of the Activity of Penicillin-Binding Proteins in Bacillus subtilis. ACS Chem Biol 15, 1242–1251 (2020). 10.1021/acschembio.9b00977

42 Zhao, G., Meier, T. I., Kahl, S. D., Gee, K. R. & Blaszczak, L. C. BOCILLIN FL, a sensitive and commercially available reagent for detection of penicillin-binding proteins. Antimicrob Agents Chemother 43, 1124–1128 (1999). 10.1128/AAC.43.5.1124

43 Bottcher, T. & Sieber, S. A. B-Lactams and b-lactones as activity-based probes in chemical biology. Med Chem Commun 3, 408 (2012). 10.1039/c2md00275b

44 Levine, S. R. & Beatty, K. E. Investigating beta-Lactam Drug Targets in Mycobacterium tuberculosis Using Chemical Probes. ACS Infect Dis 7, 461–470 (2021). 10.1021/acsinfecdis.0c00809

45 Story-Roller, E. & Lamichhane, G. Have we realized the full potential of beta-lactams for treating drug-resistant TB? IUBMB Life 70, 881–888 (2018). 10.1002/iub.1875

46 Negatu, D. A., Shin, S. J., Kim, S. Y., Jhun, B. W., Dartois, V. & Dick, T. Oral beta-Lactam Pairs for the Treatment of Mycobacterium avium Complex Pulmonary Disease. J Infect Dis 230, e241–e246 (2024). 10.1093/infdis/jiad591

47 Shin, E., Taracila, M. A., Quan, P., Khan, M. M. K., Cox, J., Patel, D., Dousa, K. M., Parmar, A., Nantongo, M., Nguyen, D. C., Rubin, E. J., Holland, S. M., Kreiswirth, B. N., Buynak, J. D. & Bonomo, R. A. Novel C5alpha-substituted carbapenems enhance Mycobacterium abscessus killing via selective target binding and reduced hydrolysis by Bla(Mab). Antimicrob Agents Chemother 69, e0017025 (2025). 10.1128/aac.00170-25

48 Story-Roller, E., Galanis, C. & Lamichhane, G. beta-Lactam Combinations That Exhibit Synergy against Mycobacteroides abscessus Clinical Isolates. Antimicrob Agents Chemother 65 (2021). 10.1128/AAC.02545-20

49 Grant, S. S., Kawate, T., Nag, P. P., Silvis, M. R., Gordon, K., Stanley, S. A., Kazyanskaya, E., Nietupski, R., Golas, A., Fitzgerald, M., Cho, S., Franzblau, S. G. & Hung, D. T. Identification of novel inhibitors of nonreplicating Mycobacterium tuberculosis using a carbon starvation model. ACS Chem Biol 8, 2224–2234 (2013). 10.1021/cb4004817

50 Tallman, K. R., Levine, S. R. & Beatty, K. E. Small-Molecule Probes Reveal Esterases with Persistent Activity in Dormant and Reactivating Mycobacterium tuberculosis. ACS Infect Dis 2, 936–944 (2016). 10.1021/acsinfecdis.6b00135

51 Hunkins, J. J., de-Moura, V. C., Eddy, J. J., Daley, C. L. & Khare, R. In vitro susceptibility patterns for rapidly growing nontuberculous mycobacteria in the United States. Diagn Microbiol Infect Dis 105, 115882 (2023). 10.1016/j.diagmicrobio.2022.115882

52 Nicklas, D. A., Maggioncalda, E. C., Story-Roller, E., Eichelman, B., Tabor, C., Serio, A. W., Keepers, T. R., Chitra, S. & Lamichhane, G. Potency of Omadacycline against Mycobacteroides abscessus Clinical Isolates In Vitro and in a Mouse Model of Pulmonary Infection. Antimicrob Agents Chemother 66, e0170421 (2022). 10.1128/AAC.01704-21

53 Garcia-Heredia, A., Pohane, A. A., Melzer, E. S., Carr, C. R., Fiolek, T. J., Rundell, S. R., Lim, H. C., Wagner, J. C., Morita, Y. S., Swarts, B. M. & Siegrist, M. S. Peptidoglycan precursor synthesis along the sidewall of pole-growing mycobacteria. Elife 7 (2018). 10.7554/eLife.37243

54 Pidgeon, S. E., Apostolos, A. J., Nelson, J. M., Shaku, M., Rimal, B., Islam, M. N., Crick, D. C., Kim, S. J., Pavelka, M. S., Kana, B. D. & Pires, M. M. L,D-Transpeptidase Specific Probe Reveals Spatial Activity of Peptidoglycan Cross-Linking. ACS Chem Biol 14, 2185–2196 (2019). 10.1021/acschembio.9b00427

55 Blum, M., Andreeva, A., Florentino, L. C., Chuguransky, S. R., Grego, T., Hobbs, E., Pinto, B. L., Orr, A., Paysan-Lafosse, T., Ponamareva, I., Salazar, G. A., Bordin, N., Bork, P., Bridge, A., Colwell, L., Gough, J., Haft, D. H., Letunic, I., Llinares-Lopez, F., Marchler-Bauer, A., Meng-Papaxanthos, L., Mi, H., Natale, D. A., Orengo, C. A., Pandurangan, A. P., Piovesan, D., Rivoire, C., Sigrist, C. J. A., Thanki, N., Thibaud-Nissen, F., Thomas, P. D., Tosatto, S. C. E., Wu, C. H. & Bateman, A. InterPro: the protein sequence classification resource in 2025. Nucleic Acids Res 53, D444–D456 (2025). 10.1093/nar/gkae1082

56 Peng, Y., Zhu, X., Gao, L., Wang, J., Liu, H., Zhu, T., Zhu, Y., Tang, X., Hu, C., Chen, X., Chen, H., Chen, Y. & Guo, A. Mycobacterium tuberculosis Rv0309 Dampens the Inflammatory Response and Enhances Mycobacterial Survival. Front Immunol 13, 829410 (2022). 10.3389/fimmu.2022.829410

57 Dousa, K. M., Shin, E., Kurz, S. G., Plummer, M., Nantongo, M., Bethel, C. R., Taracila, M. A., Nguyen, D. C., Kreiswith, B. N., Daley, C. L., Remy, K. E., Holland, S. M. & Bonomo, R. A. Synergistic effects of sulopenem in combination with cefuroxime or durlobactam against Mycobacterium abscessus. mBio 15, e0060924 (2024). 10.1128/mbio.00609-24

58 Rifat, D., Chen, L., Kreiswirth, B. N. & Nuermberger, E. L. Genome-Wide Essentiality Analysis of Mycobacterium abscessus by Saturated Transposon Mutagenesis and Deep Sequencing. mBio 12, e0104921 (2021). 10.1128/mBio.01049-21

59 Kocaoglu, O. & Carlson, E. E. Profiling of beta-lactam selectivity for penicillin-binding proteins in Escherichia coli strain DC2. Antimicrob Agents Chemother 59, 2785–2790 (2015). 10.1128/AAC.04552-14

60 The UniProt, C. UniProt: the universal protein knowledgebase. Nucleic Acids Res 45, D158–D169 (2017). 10.1093/nar/gkw1099

61 Bosch, B., DeJesus, M. A., Poulton, N. C., Zhang, W., Engelhart, C. A., Zaveri, A., Lavalette, S., Ruecker, N., Trujillo, C., Wallach, J. B., Li, S., Ehrt, S., Chait, B. T., Schnappinger, D. & Rock, J. M. Genome-wide gene expression tuning reveals diverse vulnerabilities of M. tuberculosis. Cell 184, 4579–4592 e4524 (2021). 10.1016/j.cell.2021.06.033

62 Kang, C. M., Nyayapathy, S., Lee, J. Y., Suh, J. W. & Husson, R. N. Wag31, a homologue of the cell division protein DivIVA, regulates growth, morphology and polar cell wall synthesis in mycobacteria. Microbiology (Reading) 154, 725–735 (2008). 10.1099/mic.0.2007/014076-0

63 Melzer, E. S., Sein, C. E., Chambers, J. J. & Siegrist, M. S. DivIVA concentrates mycobacterial cell envelope assembly for initiation and stabilization of polar growth. Cytoskeleton (Hoboken) 75, 498–507 (2018). 10.1002/cm.21490

64 Mukherjee, P., Sureka, K., Datta, P., Hossain, T., Barik, S., Das, K. P., Kundu, M. & Basu, J. Novel role of Wag31 in protection of mycobacteria under oxidative stress. Mol Microbiol 73, 103–119 (2009). 10.1111/j.1365-2958.2009.06750.x

65 Halbedel, S. & Lewis, R. J. Structural basis for interaction of DivIVA/GpsB proteins with their ligands. Mol Microbiol 111, 1404–1415 (2019). 10.1111/mmi.14244

66 Meniche, X., Otten, R., Siegrist, M. S., Baer, C. E., Murphy, K. C., Bertozzi, C. R. & Sassetti, C. M. Subpolar addition of new cell wall is directed by DivIVA in mycobacteria. Proc Natl Acad Sci U S A 111, E3243–3251 (2014). 10.1073/pnas.1402158111

67 Jiao, W., Blackmore, N. J., Nazmi, A. R. & Parker, E. J. Quaternary structure is an essential component that contributes to the sophisticated allosteric regulation mechanism in a key enzyme from Mycobacterium tuberculosis. PLoS One 12, e0180052 (2017). 10.1371/journal.pone.0180052

68 Webby, C. J., Jiao, W., Hutton, R. D., Blackmore, N. J., Baker, H. M., Baker, E. N., Jameson, G. B. & Parker, E. J. Synergistic allostery, a sophisticated regulatory network for the control of aromatic amino acid biosynthesis in Mycobacterium tuberculosis. J Biol Chem 285, 30567–30576 (2010). 10.1074/jbc.M110.111856

69 Blackmore, N. J., Reichau, S., Jiao, W., Hutton, R. D., Baker, E. N., Jameson, G. B. & Parker, E. J. Three sites and you are out: ternary synergistic allostery controls aromatic amino acid biosynthesis in Mycobacterium tuberculosis. J Mol Biol 425, 1582–1592 (2013). 10.1016/j.jmb.2012.12.019

70 Betts, J. C., Lukey, P. T., Robb, L. C., McAdam, R. A. & Duncan, K. Evaluation of a nutrient starvation model of Mycobacterium tuberculosis persistence by gene and protein expression profiling. Mol Microbiol 43, 717–731 (2002). 10.1046/j.1365-2958.2002.02779.x

71 Shrivastava, T. & Ramachandran, R. Mechanistic insights from the crystal structures of a feast/famine regulatory protein from Mycobacterium tuberculosis H37Rv. Nucleic Acids Res 35, 7324–7335 (2007). 10.1093/nar/gkm850

